# *Sericanthe etugei* and *S. onanae* (Coffeeae Rubiaceae) new cloud forest tree species Endangered in Southwest Cameroon & Bioko (Equatorial Guinea)

**DOI:** 10.1101/2025.02.24.639651

**Authors:** Martin Cheek, Jean Michel Onana, Bonaventure Sonké

## Abstract

Two new species of cloudforest tree are described and illustrated from the Cross-Sanaga Interval within the Lower Guinea Domain. *Sericanthe etugei* Cheek from South West and Littoral Regions of Cameroon is species from the Cameroon Highlands, and *Sericanthe onanae* Cheek from South West Region, Cameroon, and Bioko (Equatorial Guinea).

*Sericanthe etugei* is similar to *Sericanthe leonardii* (N. Hallé) Robbr. of DRC (Kivu) and Rwanda, differing in the stipular awn 0.25 mm long (vs 1 – 4 mm), the adaxial leaf midrib and secondary nerves glabrous (vs hairy), the tertiary nerves conspicuously scalariform with the naked eye (vs not visible). The species is also similar to *Sericanthe testui* var. *testui,* differing in being a tree (vs liana), with stems not decorticating (vs decorticating), the calyx truncate in bud (vs closed) and in the pubescent ovary (vs glabrous).

*Sericanthe onanae* is similar to *S. mpassa* Sonké & Robbr. of lowland forest in Gabon, differing from that species in that the leaves have an acumen (0.5 – )0.7 – 0.8(– 1.2) cm long vs 0.3 – 0.5 cm, secondary nerves are 7 – 10 (– 12) on each side of the midrib (vs 4 – 6 in *S. mpassa*), and that the stipules are 5 – 6.5( – 7) × 3 – 5(– 6) mm ( vs 2.5 mm long) and truncate (vs overtopped by an awn c. 1 mm long). It is also similar to *S. jacfelicis* (N. Halle) Robbr., differing in that the leaves are smaller (<15 cm long vs 16 – 20 cm long), the abaxial surface of the leaf blade has 7 – 10( –12) lateral nerves on each side of the midrib (vs 5 –7) and has a glossy surface and papery texture (vs matt, coriaceous), the petioles are hairy (vs glabrous), the stipules are 5 – 6.5( – 7) mm long, truncate, lacking an awn or arista, (vs 3 –4 mm long, with an arista 0.5 – 1 mm long). These two new species are both provisionally assessed as Endangered EN B2ab(iii) with the 2012 IUCN standard.

The new species are briefly discussed in the context of other newly discovered submontane species from the Cross-Sanaga Interval. We speculate that the seeds of these species may be primate-dispersed.

## Introduction

### The genus *Sericanthe*

*Sericanthe* is placed in the tribe Coffeeae as defined by Davis *et al*. (2007). The tribe is characterised by the paired axillary 1 or few-flowered inflorescences, each flower is subtended by 1 – 4-cupular calyculi and has flowers with a bilobed stigma. *Sericanthe* is characterised by the white, adpressed, silky hairs of the flowers (giving rise to the generic name), the basifixed, unappendaged stamens generally with short flat filaments, the often 6 – 8-merous flowers and the presence in most species of bacterial bodies (known as galls or nodules) in the leaves, as either lines along the midrib or dots and dashes throughout the blade. *Sericanthe* is the only genus of the tribe with these bacterial bodies, although they also occur in three other genera of the family (Cheek & Onana 2024). The genus also has large (c. 1 cm diam.), entire seeds with a broad hilum and associated placental fleshy tissue that can cover a third or more of the seed surface, and with a small, transverse embryo within the large, hard endosperm. The pollen has a verrucate surface unlike the reticulate state of the genus *Tricalysia* A.DC in which it was formerly included (Robbrecht 1978).

*Coffea* L. is sister to the remainder of the Coffeeae tribe, with the next diverging branch the *Argocoffeopsis* clade (Cheek *et al*. 2018a). *Sericanthe* is placed in a clade with *Empogona*, *Discospermum* Dalzell, and *Diplospora* DC. in Tosh *et al*. (2009). However, in Cheek *et al*. (2018a) *Sericanthe* is the next diverging branch after the *Argocoffeopsis* clade and sister to the rest of the tribe. Arriola *et al*. (2018) included additional Asian representatives in their sampling and postulated that *Tricalysia sensu stricto* is sister to the clade (*Sericanthe* (*Diplospora*, *Empogona*)). A key to the 12 genera in tribe *Coffeeae* can be found in Cheek *et al*. (2018a). This tribe has its highest generic diversity in Cameroon with nine of the 12 genera present (Cheek *et al*. (2018a).

*Sericanthe* was erected by Robbrecht (1978) to replace the name *Neorosea* N. Hallé (1970, 1972) since the last author had by error typified his generic name on a species of *Tricalysia* A.DC instead of a species of the genus that he was the first to recognise as distinct from *Tricalysia*. Since the time of Robbrecht’s foundational revision of the genus, two more species were transferred to it from *Tricalysia* and accommodated in the newly erected subgenus *Macrocarpa* Robbr. (Robbrecht 1981). New subspecies of *S. andongensis* (Hiern)Robbr. were recognised by Bridson in Bridson & Verdcourt (2003) and by Jordaan & Steyn (2012). Four new species were added by Sonké *et al*. (2012) in a revision of the Lower Guinea species. Finally, a new species was published from the Chimanimani Mts of the Zimbabwe/Mozambique border by Wursten *et al*. (2020). Currently 22 species are recognised (POWO, continuously updated). Five species were recognised for Cameroon (Onana 2011) and four for Gabon (Sosef *et al*. 2006) but these numbers are increased by Sonké *et al*. (2012).

The genus extends from Senegal in West Africa to Tanzania in the east and ranges as far south as northern South Africa (Robbrecht 1978). It is not recorded from Uganda, Kenya nor NE Africa (Ethiopia, Somalia, S. Sudan). The species are deciduous or evergreen shrubs or small to medium size trees of woodland or forest.

Many forest species of the genus have small ranges and so are especially at risk from habitat destruction. Four species appear as threatened in the Red Data Book of Cameroon Plants (Onana & Cheek 2011) but only one of these (Cheek & Lovell 2021) appears among the 12 species currently listed as globally threatened on iucnredlist.org (accessed Jan. 2025).

### Botanical surveys in the Cross Sanaga Interval

The new species published in this paper resulted from the long-term survey of plants in Cameroon to support improved conservation management led by botanists from Royal Botanic Gardens, Kew, the IRAD (Institute for Research in Agronomic Development)– National Herbarium of Cameroon, Yaoundé and University of Yaoundé I. This study has initially focussed on the Cross-Sanaga interval (Cheek *et al*. 2001) which contains the area with the highest species and generic diversity per degree square in tropical Africa (Barthlott *et al*. (1996); Dagallier *et al*. (2020). To date, three new genera have been discovered through these studies, *Kupea* Cheek & S.A. Williams (Triuridaceae, Cheek *et al*. 2003), *Korupodendron* Litt & Cheek (Vochysiaceae, Litt & Cheek 2002) and *Kupeantha* Cheek (Rubiaceae, Cheek *et al*. 2018a) and hundreds of new species including Achoundong & Cheek (2003); Alvarez-Aguirre *et al*. (2021); Cheek (2000; 2002); Cheek *et al*. (2018b; 2021); Cheek & Bridson (2002); Couvreur *et al*. (2022); Ghogue *et al*. (2017); Gosline & Malécot (2011); Gosline *et al*. (2014; 2022); Lachenaud (2019); Lachenaud *et al*. (2013); Mackinder *et al*. (2010); Mackinder & Pennington (2011); Prance & Jongkind (2015); Salazar *et al*.(2002); Sonké (2000); Sonké *et al*.(2002a; 2002b); Stone & Cheek (2018); Stone *et al*. (2008) and van der Burgt *et al*. (2015).

The herbarium specimens collected in these surveys formed the primary data for the series of Conservation Checklists that began at Mt Cameroon (Cheek *et al*. 1996). Two new national parks have resulted from these checklists so far, the Mt Cameroon National Park, and the Bakossi Mts National Park, the first to be specifically for plant conservation (rather than for animals) in Cameroon.

Each of these conservation checklists has a section on Red Data species. Building on these, a national Red Data Book of the Plants of Cameroon was delivered (Onana & Cheek 2011), the first for any tropical African country. The Red Data book in turn helped the Tropical Important Plant Areas programme (Darbyshire *et al*. 2017) that resulted in the book on the Important Plant Areas of Cameroon that evidenced and mapped priority areas for plant conservation in Cameroon with local partners (Murphy *et al*. 2023).

In this paper we formally publish two species of cloud forest (submontane) tree of the genus *Sericanthe* that had previously been described under the informal names *S.* sp. A and *S.* sp. B. (Cheek in Cheek et al 2004). These are here named as *Sericanthe etugei* and *Sericanthe onanae* respectively. This will facilitate IUCN accepting conservation assessments for the species, so that these can be incorporated into conservation prioritisation initiatives.

*Sericanthe* sp A, because of the linear bacterial nodules, the montane habitat, the abaxial leaf indumentum being on the nerves only, the style and ovary being pubescent, the calyx open in bud with truncate margin, would key out in Robbrecht (1978) to *Sericanthe leonardii* Robbr. of DRC-Kivu and Rwanda. In the key of Sonké et al. (2012), because the leaf base is cuneate, and the intersecondary (tertiary) nerves are very apparent, forming a dense pattern of parallel veinlets perpendicular to the midvein, the species would instead key out as *S. ‘rumpi’*.

However, in the last species the veinlets are even more apparent, and more numerous than in *Sericanthe* sp A, and in *S. ‘rumpi’* the abaxial leaf surface is glossy not dull, and domatia and bacterial nodules are absent, in contrast to *Sericanthe* sp A.

*Sericanthe* sp B, since it has absent or difficult to observe bacterial nodules, a montane forest habitat, leaf blade only sparsely hairy beneath (on the veins) and young twigs glabrous, keys out in Robbrecht (1978) to couplet 13 where it most nearly fits *S. jacfelicis* (N. Halle) Robbr., differing in that the leaves are much smaller (<15 cm long vs 16 – 20 cm long) than in *S. jacfelicis. Sericanthe* sp. B further differs from *S. jacfelicis* in that the abaxial surface of the leaf blade has 7 – 10( –12) lateral nerves on each side of the midrib (vs 5 –7) and has a glossy surface and papery texture (vs matt, coriaceous), the petioles are hairy (vs glabrous), the stipules are 5 – 6.5( – 7) mm long, truncate, lacking an awn or arista, (vs 3 –4 mm long, with an arista 0.5 – 1 mm long) and the flowers are borne on older stems, several nodes beneath the leaves (vs among the leaves on young stems). A remarkable feature of *Sericanthe* sp. B is that the exterior of the of corolla tubes have longitudinal bands which are glabrous, alternating with those that are white hairy, unique in the genus.

According to Sonké et al. (2012), because the bacterial nodules are inconspicuous or absent, the leaf base is cuneate, the abaxial surface inconspicuously hairy, the tertiary nerves (inter secondaries) not conspicuously parallel, and because of the leaf blade and petiole dimensions, *Sericanthe* sp B keys out to couplet 11, and most closely fits *S. mpassa* Sonké & Robbr. of lowland forest in Gabon. However, it differs from that species in that the number of secondary nerves on each side of the midrib are 7 – 10 (– 12) (vs 4 – 6 in *S. mpassa*), and that the stipules are much longer than those of this species (5 – 6.5(– 7) mm vs 2.5 mm long) and truncate (vs overtopped by an awn c. 1 mm long).

## Material and methods

The methodology for the surveys in which this species was discovered is recorded in Cheek & Cable (1997). Nomenclatural changes were made according to the Code (Turland *et al*. 2018). Names of species and authors follow IPNI (continuously updated). Herbarium material was examined with a Leica Wild M8 dissecting binocular microscope fitted with an eyepiece graticule measuring in units of 0.025 mm at maximum magnification. The descriptions are based on herbarium specimens, spirit material and pictures when available, and data derived from field notes. The drawing was made with the same equipment with a Leica 308700 camera lucida attachment. Specimens were inspected from the following herbaria: BM, BR, K, P, WAG, YA using the conventions of Davies *et al*. (2023) and specimens were also viewed on GBIF (continuously updated), and species records checked on POWO (continuously updated). The format of the description follows those in other papers describing new species in Coffeeae e.g. Cheek *et al*. (2018a). Terminology follows Beentje & Cheek (2003), but for specialised structures of Rubiaceae, e.g. for stipule colleters, and domatia, generally follows Robbrecht (1987; 1988). Phytogeographical considerations follow White (1979; 1983; 1993) but with his chorological categories (“regional (sub)centre of endemism” and other) simplified into Region and Domain. All specimens cited have been seen unless indicated “n.v.” The conservation status of the new species was assessed according to IUCN Red List Categories and Criteria, version 3.1 (IUCN 2012; IUCN Standards and Petitions Subcommittee 2015). The extent of occurrence and the area of occupancy were estimated with GeoCAT (http://geocat.kew.org) with a grid size of 2 km x 2 km. Herbarium codes follow Index Herbariorum (Thiers continuously updated).

## Taxonomic results

***Sericanthe etugei*** Cheek *sp. nov*.

Type: Cameroon, South West Region, Korup National Park, Path from Ekundu Kundu to top of Mt Juahan, 900 m, fl. fr. 27 April 1996, *Cheek* 8235 with Elangwe, Etuge, Nana, Satabie, Gosline (K holo. K001900901; iso. MO, YA).

*Sericanthe* sp. A Cheek (2004: 391).

*Sericanthe testui* (Hallé) Robbr. var *testui* sensu Sonké et al. (2012: 548) quoad *Etuge & Thomas* 328 and *Achoundong* 1232 non sensu Hallé (1970)

Differing from *Sericanthe leonardii* (N. Hallé) Robbr. of DRC-Kivu and Rwanda in the stipular awn 0.25 mm long (vs 1 – 4 mm), the adaxial leaf midrib and secondary nerves glabrous (vs hairy), the tertiary nerves conspicuously scalariform with the naked eye (vs not visibly scalariform), the stem hairs tightly appressed, straight, c. 0.6 mm long (vs spreading, c. 0.25 mm long), the calyx sparsely puberulent, c. 40% covered in minute hairs (vs completely covered in long sericeous hairs), the corolla apex in bud rounded (vs apiculate), the ovary-hypanthium concealed by the calyculi (vs exposed, visible). For differences with *Sericanthe testui* see Table 1, below.

**Table 1.** Characters separating *Sericanthe testui* from *Sericanthe etugei sp.nov.* Data for the first species taken from (Hallé 1970) and Robbrecht (1981) and five herbarium specimens at K.

|  | *Sericanthe testui* | *Sericanthe etugei* *sp. nov.* |
| --- | --- | --- |
| Altitudinal range (m) and habitat | 50 – 800. Lowland rain forest and swamps | (700 –) 900 – 1750. Slopes in submontane forest |
| Habit | Liana or climbing shrub | Tree |
| Epidermis conspicuously decortivating in scales | Yes | No |
| Petiole length | 4 – 8 | (4 – )6 – 12( – 14) |
| Apiculus of acumen | Absent or indistinct | Distinct, 0.2 x 0.1 mm |
| Pit domatia presence | Absent | Present (sometimes absent). |
| Calyx in flower bud | Closed, concealing corolla | Truncate, corolla exposed |
| Calyx fissures at anthesis | Fissured ± to the calyx base | Fissures ± 1/4 length to base |
| Corolla tube: lobe lengths | Tube > lobes | Lobes > tube |
| Corolla lobe length (mm) | 3 – 4.5 | 8.5 – 9.5 |
| Diam. corolla at anthesis (mm) | 4 – 9 | 12 – 14(–18) |
| Staminal filament length (mm) | ±1 | 1.5 – 2 |
| Ovary surface | Glabrous | Densely pubescent |
| Fruit surface | Non-verrucate, glabrous | Verrucate, hairs 0.2 mm long |
| Geographic Range | South and Littoral Regions Cameroon, Gabon, and bas-Congo (DRC). | South West and Littoral Regions Cameroon |

<u>Evergreen tree</u> (2 – )6 – 10 m tall, trunk white, slender (*Cheek* 8285). Leafy stems 14 – 32 cm long, rarely branched, internodes at first pale brown, terete, (1 – )1.7 – 3.2(– 3.8) cm long, 1.5 – 3.2 mm diam., (4 – )5 – 8( – 12) pairs of leaves present per stem, leaves of a pair subequal in shape and size; indumentum of stems (extending to the petioles, abaxial midrib, secondary nerves, abaxial stipule) of dense (c. 80 – 100 % cover of stem, petioles, abaxial midrib) appressed, simple, white or pale yellow-brown stiff, straight hairs 0.25 – 0.6 mm long, persisting until the 8^th^ internode or more, epidermis then becoming glabrous, dull brown, red or pale grey, with few longitudinal rounded ridges, lenticel-free, not decorticating. <u>Stipules</u> long-persistent, brown, membranous, subcylindrical, 4 – 6 × 2 – 3.5 mm completely sheathing (limbs ±absent at the distal 3– 4 internodes or limbs rounded, shallow, c. 1 x 1.8 mm), midrib ridged distally for c.1.5 mm long (Fig. 1C) or extending the full length of stipule, sometimes extending as a short awn 0.25 (– 0.5) mm long; stipules of older internodes (Fig. 1C&D) splitting nearly to the base and awn appearing more prominent; outer surface with indumentum as stem but sparse (c. 50% cover), inner surface glabrous apart from the basal c. 0.5 mm with sparse silky appressed hairs c. 2.5 mm long; colleters winged (Robbrecht 1987a), hairy and very sparse and inconspicuous, pale dull red, c. 0.4 x 0.1 mm. <u>Leaf-blades</u> papery, upper surface drying dark brown to black, lower surface drying bright red brown (distal leaf pair) fading somewhat to duller brown (leaves of subsequent internodes), the abaxial midrib bright white (indumentum of young leaves) or pale brown, oblanceolate, rarely oblanceolate-oblong, (6.2 – )9 – 11( – 12.5) × (2 – )3 – 4 (– 4.5) cm, apex acuminate, acumen 0.2 – 1.25 cm, apiculus of acumen cylindrical, 0.2 x 0.1 mm, clothed in appressed white hairs, base of blade cuneate to acute, rarely nearly obtuse, shortly decurrent down the distal petiole; adaxial surface with midrib sunken and hairy proximally otherwise glabrous and nerves flat and inconspicuous; abaxial surface with midrib raised, brochidodromous (secondary nerves seemingly looping); domatia conspicuous, both tuft and shallow pit types (Robbrecht 1988), c. 0.7 mm diam., the pits glossy, dark brown, glabrous surrounded by a few erect hairs c.0.3 mm long, or the hairs more numerous, 40 – 50, obscuring the pits (which may be absent in some specimens); bacterial galls/nodules very inconspicuous (due to the indumentum) and seemingly not always present, black, linear 1 – 1.5 cm long, longitudinal along the flanks of the proximal midrib abaxially; secondary nerves (4 – )5 – 6(– 7) on each side, arising at 45 – 70 degrees from the midrib, curving upwards and gradually joining the nerve above through a branch, forming a weakly looping infra-marginal nerve; and secondary nerves on abaxial surfaces with indumentum as stem, hairs slightly sparser and shorter (c. 30 – 40% cover, 0.1 –0.3 mm long), otherwise glabrous on both surfaces; tertiary nerves raised, concolorous, moderately conspicuous, scalariform, patent to midrib, 14 –18 between each pair of secondary nerves (8 –10 towards leaf apex), sparsely branched, glabrous; quaternary nerves inconspicuous; margin slightly recurved and thickened, glabrous. <u>Petiole</u> patent, shallowly canaliculate (4 – )6 – 12( – 14) × 0.9 – 1.1 mm, indumentum as stem. <u>Inflorescences</u> slightly supra-axillary, inserted c.1 mm above the axil, single (Fig. 1E) or in both axils of a node (very rarely a second superposed inflorescence per axil), flowering nodes per stem 1 – 2(–4), occurring at the 5 – 8 th node from the apex, flowers 1 per inflorescence, inflorescence highly condensed; calyculi 2 – 3 per flower, papery-translucent, ±cup-shaped, completely concealing the white pubescent axis in flower (Fig. 1E), but visible in fruit (see below), indumentum of outer surfaces ±sparsely white shortly sericeous, hairs as stem, inner surfaces glabrous, colleters not observed. 1^st^ order (proximal) calyculus brown, papery, cupular, 1 – 1.2 × 1.75 – 2.5 mm, subtending 2^nd^ order calyculi (if present), margin usually truncate, indumentum sparse, 50% cover, hairs short, colourless, variously oriented, 0.2 – 0.25 mm long. 2^nd^ order (medial) calyculus if present similar to 1st order calyculus but slightly larger. 3^rd^ order calyculi, sparsely and shortly white-sericeous, cylindrical to subcampanulate, 1.75 – 2.5 x 2.5 – 3.75 mm, shortest and widest when fissured, usually with two long awns c. 1 mm long, and two short awns c. 0.25 mm long. <u>Flowers</u> hermaphrodite, homostylous, 6 –7-merous, 12 – 14(– 18) mm diam. c. 13 mm long. Ovary-hypanthium subcylindric, c. 2.5 × 0.75 mm, concealed within 3^rd^ order calyculus. Densely white hairy, hairs 0.25 mm long. Calyx cylindrical, not enclosing corolla in bud, indumentum as calyculi, but outer surface with only c. 40% cover, inner surface with a few hairs at base, colleters 0.25 x 0.02 mm; calyx tube (3 – )3.7 – 4 x 4 – 4.5 mm; at anthesis with 2 or 3 short longitudinal fissures c. 1 mm long (Fig. 1F); calyx lobes shallow, 6 –7, triangular or rounded, with apiculus, erect, 0.2 × 1 – 1.1 mm. Corolla apex exposed above calyx in bud, conical, apex rounded; at anthesis white (*Cheek* 8235), drying black, tube subcylindrical, 4.5 – 5(– 6.5) × 2 mm, widest at mouth, except for the glabrous proximal 1 mm, shortly white pubescent outside, hairs 0.1 – 0.5 mm long; inner surface with proximal 2.5 – 3 mm glabrous, hair band distal, 2 mm long, with crinkled flat white hairs 0.1 – 0.6 mm long, distal 1 mm with hairs longest and densest (70% cover) and exserted, otherwise 40% cover;Corolla lobes 6 –7, left-contorted in bud, spreading to slightly reflexed at anthesis and longer than tube (Fig 1F), elliptic-oblong, 8.5 – 9.5 × 1.7 – 2.8 mm, apex obtuse-rounded, adaxially glabrous, abaxially white puberulent, hairs c. 0.1 mm long, appressed; Stamens 6 – 7, anthers fully exserted, glabrous, anthers oblong, 2.25 × 0.4 – 0.7 mm, connective conspicuous the entire length, drying dark brown, basifixed, apical connective appendage absent; filaments 1.5 – 2 x 0.3 – 0.5 mm, inserted at mouth of tube, slightly dorsiventrally flattened. Disc subcylindrical, 0.35 × 0.5 mm, glabrous. Style 5.5 – 6 × 0.2 – 0.3 mm (narrowest at base, widest at apex), exserted slightly beyond the anthers, glabrous at apex, otherwise with patent white hairs c. 0.25 mm long (Fig. 1E); style arms 2, opposed, forming a T-shape, each ellipsoid, 1.4 – 1.6 x 0.7 mm, stigmatic surfaces smooth (Fig. 1G). Ovary 2– locular, ovules 2 per locule, partly immersed in the swollen placentae. Fruit orange (dark red in *Cheek* 8235) when mature, subglobose or shortly ellipsoid, 12 – 13 (– 23) x 7 – 8.5(– 25) mm, apex crowned by the persistent calyx (Fig. 1H), calyx tube thickened, lobes papery, brown; disc accrescent. 32 mm wide (Fig 1L); equator often with a raised ridge (Fig. 1J); base covered with the appressed 3^rd^ order calyculus; peduncle slightly accrescent (3 –)5 – 6 x 1.5 mm, naked apart from persistent calyculi(-us), surface verrucate, projections low, rounded, 0.2-0.3 mm diam., hairs sparse, covering c.10 % of surface, suberect, 0.2 mm long (Fig. 1K); fruit wall fleshy, about 1 mm thick (rehydrated), inner wall glabrous, endocarp parchment-like, 1 – 4-seeded. Seeds subglobose (1-seeded fruit) c. 10 x 12 x 8 mm, or hemispherical or segment-shaped, 9 – 10.5 x 7 – 9.5 x 5.5 – 9 mm, testa glossy dark brown, hilum white, covering 40 – 50% of surface, endosperm bright white, embryo transverse, cylindric-conical c. 1.5 mm long.

**Fig. 1.**
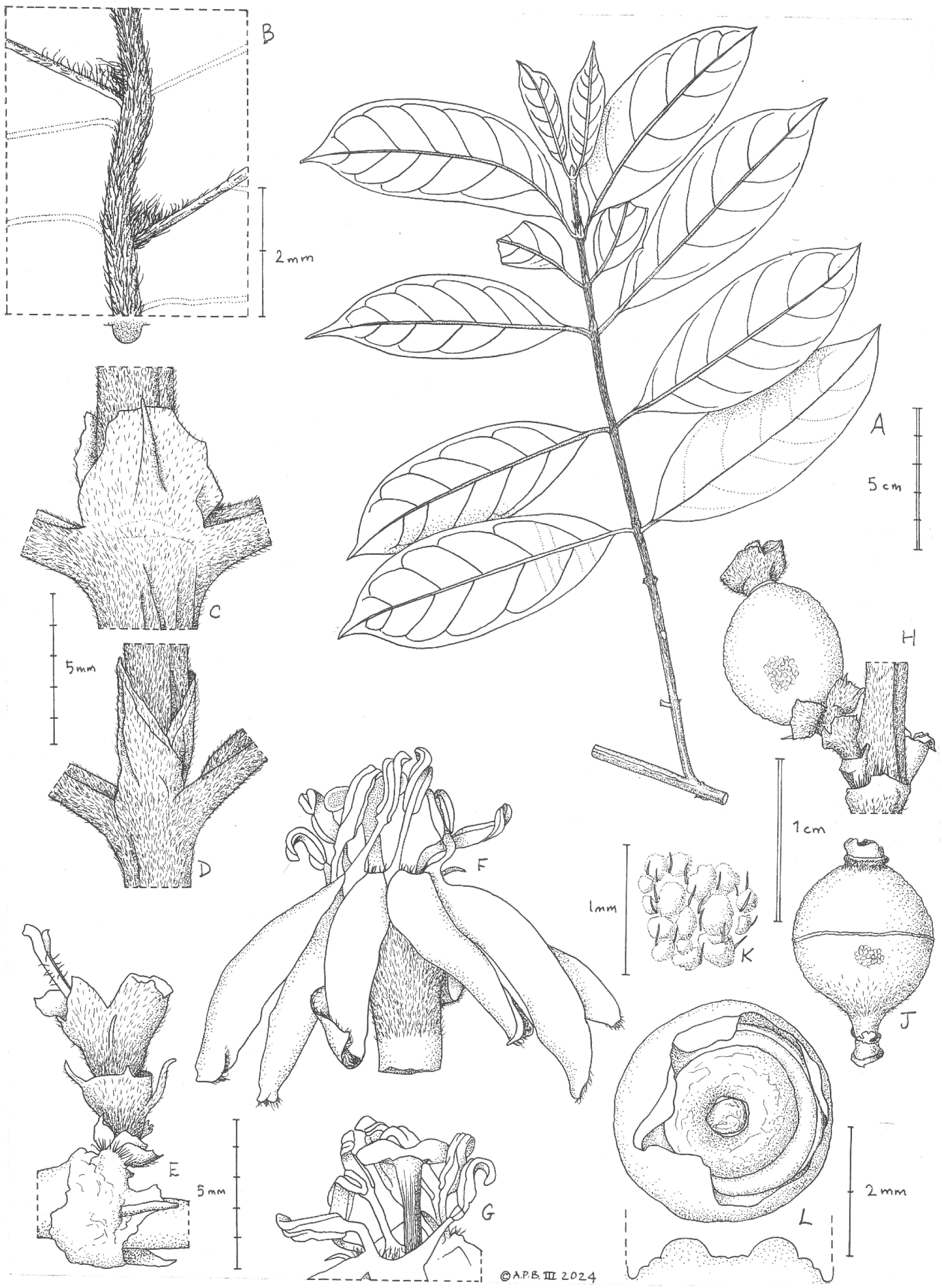
*Sericanthe etugei* **A** habit; **B** leaf abaxial showing two domatia; **C & D** fissured stipules from old internodes; **E** inflorescence (corolla fallen); **F** flower, detached from calyx-hypanthium showing corolla, stamens and style, side view; **G** part of F from rear to show style and stigma; **H** young fruit in situ; **J** detail of surface of fruit; **K** another fruit showing an equatorial line; **L** apical view of fruit to show accrescent disc. **A-D, H** from *Cheek* 8925 (K); **E** from *Etuge & Thomas* 328 (K); **F&G** from *Cheek* 8235; **J-L** from *Thomas* 10550 (K). Drawn by ANDREW BROWN.

**SPECIMENS EXAMINED. CAMEROON. SOUTH WEST REGION. Korup National Park**, Path from Ekundu Kundu to top of Mt Juahan, 900 m, fl. fr. 27 April 1996, *Cheek* 8235 with Elangwe, Etuge, Nana, Satabie, Gosline (K holo. K001900901; iso. MO, YA); Path from forest camp (near Ekundu Kundu) to top of Mt Juahan (200 m base, c. 1100 m at top in alt.). Top of mountain, fr. 30 April 1996, *Cheek* 8285 with Balcha, Joseph, Kennedy, Etuge, Bodier (K, MO, YA); **Nta Ali Forest Reserve**, fl. buds, 10 June 1983, *Achoundong* 1232 (BR image!, YA n.v.); 800 – 1250 m elev., fr. 12 March 1995, *D.W. Thomas* 10550 (K, YA); **Bakossi Mts**. Path N.W. Elumseh-Mejelet, Bakossi, Bangem, 1750 m. fl. fr. 6 Oct. 1986, *Etuge & D.W.Thomas* 328 (GH, K, MO n.v., YA); Kodmin, 0.5 km on road to Ndip, 1500 m, fr. 18 Jan. 1998, *Cheek* 8925 with Gosline, Kemei, Taylor, Abwe Gabriel, Frith (BR, K, MO, SCA, YA); **Mt Kupe**, Path on to ridge between Daniel’s saprophyte site and Kupe rock, 1000 m, fr. 30 May 1996, *Cable* 2731 with Amy Sui, Satabie, Ajang D. (BR, K, MO, SCA, YA); Path from Kupe village to lohe path to summit, near junction, fl.buds, fr. 17 July 1996, *Cable* 3869 with Bishop P., Ajang D., Ndue H., Muellner L. (BR, K, MO, SCA, YA).

**LITTORAL REGION. Ebo Forest.** Bekob, submontane forest after the end of the transect leading to the North, 830 m, fr. 17 Feb. 2006, *Tchiengue* 2517 with Oben, B., Ekwoge Enang Abwe (K, YA).

**HABITAT & ECOLOGY.** Submontane (cloud) forest with *Santiria trimera*, *Syzygium staudtii, Leonardoxa africana* (*Thomas* 10550) *Garcinia* spp., *Allanblackia gabonensis* and *Zenkerella citrina* (*Tchiengue* 2715); (700 –)900 – 1500(– 1750) m alt.

DISTRIBUTION. Cameroon. Endemic

**ETYMOLOGY.** Named in honour (meaning ‘Etuge’s Silky Flower) of the late Martin Etuge Ekwoge (-2020) of Bakossi, known professionally as Martin Etuge, botanical specimen collector from 1984 – 2000. This is the sixth species known to be named for him, the other species being *Kupea martinetugei* Cheek & S.A. Williams (Triuridaceae), *Psychotria martinetugei* Cheek (Rubiaceae), *Cola etugei* Cheek (Sterculiaceae s.s.), *Uvariopsis etugeana* Couvreur & Dagallier (Annonaceae), *Impatiens etugei* Cheek (Balsaminaceae). For further notes on Martin Etuge see Cheek et al. (2023a).

**CONSERVATION STATUS.** At four of the eight known sites for this species, equating to three locations (near Bangem, Kodmin and Kupe settlements), there are threats of habitat clearance, mainly from small-holder agriculture from nearby villages. However, at the Ebo site, industrial logging is the imminent threat, likely to be followed by agriculture. Only at the Mt Juahan sites, which are inside Korup National Park, is the species formally protected, yet at Nta Ali, the northernmost location, because of the steep slopes, there also the species may be secure from habitat clearance. We consider there to be five threat-based locations for the species.

Using the IUCN required 2 km x 2 km grid cells we calculate the area of occupancy as 32 km^2^. The extent of occurrence was calculated as 7353 km^2^ which qualifies for the Vulnerable level of threat. However, the AOO, together with the number of locations and threats stated, merit a provisional assessment of Endangered EN B2ab(iii) for *Sericanthe etugei*.

It is possible that the species may be found at other sites, but numerous surveys have already been conducted in the range of this species and in surrounding areas: (Cheek *et al*. 1992; Cheek *et al*. 1996; Cable & Cheek 1998; Cheek *et al*. 2000; Maisels *et al*. 2000; Chapman & Chapman 2001; Cheek *et al*. 2004; Harvey *et al*. 2004; Cheek *et al*. 2006; Cheek *et al*. 2010; Harvey *et al*. 2010; Cheek *et al*. 2011).

Conservation actions to help protect this species should start with a sensitisation programme with the local communities concerned to enable understanding of the importance of conservation of this species and to support species identification and protection. Development of an action plan to protect the species is recommended. Since seed seems to be abundantly produced (most of the specimens have fruits) propagation from seed in local nurseries seems a good possibility.

**NOTES**. We found that the very first gathering of the novelty goes back to 1983, when Achoundong collected a specimen with only flower buds in the Nta Ali area. The collection of *Sericanthe etugei* first known to MC was *Etuge & D.W. Thomas* 328 from Elumseh-Mejelet in the Bakossi Mts. It was determined as a *Sericanthe* by Bridson in 1988. Three more collections were made in 1995 and 1996 from the Bakossi Mts and Mt Kupe and the taxon was identified as new species, S. sp A. in Cheek et al. (2004). Two specimens collected from Mt Yuhan in Korup in 1996 and another collected from Nta Ali in 1995, were not identified as S. sp. A until later. These include the only specimen with flowers at anthesis, selected as type for this reason.

*Etuge & D.W. Thomas* 328 was referred to *Sericanthe testui* (Hallé) Robbr. var. *testui* with caveats, in Sonké et al. (2012), without reference to Cheek et al. (2004). A second collection referred to in the same way in Sonké et al. (2012) is *Achoundong* 1232 from Nta Ali with only flower buds, yet the image of the BR sheet leaves no doubt that it is also *Sericanthe etugei*. The caveats in Sonké et al. (2012) concerning these two specimens being identified as *Sericanthe testui* relate to their domatia and stem hairness “the domatia are weakly developed; this material can be confused with *S. rabia*”. With more and much more complete material, these caveats seem prescient.

*Sericanthe testui* and *Sericanthe etugei* are indeed similar and may even be sister to each other. However, despite the many similarities, they are separated by numerous traits including many which are qualitative (Table 1). It is possible that the similarities are due to convergent evolution. For the moment, *Sericanthe testui* is not known from South West Region, Cameroon, the main range of *Sericanthe etugei*, but only to the south and east (see Table 1). The two species are known to be sympatric only at one location, although in different altitudinal zones, in the Ebo Forest of Littoral Region which is both the current eastern limit of *Sericanthe etugei*, and the western limit of *Sericanthe testui*.

That *Sericanthe testui* is a liana is not universally clear from the notes of the specimens referred to by Hallé (1970) in the protologue, because little metadata was recorded by Le Testu and by Bates, the main collectors of the original material. However, *Le Testu* 9296 states “liana”. Yet, more recent collections of the species e.g. *Osborne* 64 (“Liana 10 m+” Ebo forest) leave no doubt that this taxon is a liana in habit (see also in this respect *Osborne* 195, *J.J.F.E. de Wilde et al.* 334, *J.M. & B.Reitsma* 898, *Breteler* 6923). The slender winding stems seen in those specimens where habit is not indicated suggest that these are also climbers, in contrast with all specimens of *S. etugei* which are stated by collectors to be trees, and which have straight, stout leafy stems consistent with this habit.

*Sericanthe etugei* also has similarities with *S. leonardii* of DRC-Kivu and Rwanda, in fact one specimen of the first, *Tchiengue* 1275 had been identified and incorporated as *S. leonardii* at K despite the large geographical separation. Differences are given in the diagnosis.

*Sericanthe onanae* Cheek *sp. nov*.

Type: Cameroon, South West Region, Banyang Mbo Wildlife Reserve, Ebamut, 1200 m alt., fl. buds, fr. 8 Nov. 2001, *Onana* 1949 with Nana, Ghogue, Etuge Ambros, Okong, Kongor, Fobia (holotype K barcode K001900938; isotypes BR, MO, YA).

*Sericanthe* sp. B Cheek (2004: 391).

Most similar to *S. mpassa* Sonké & Robbr. of lowland forest in Gabon, differing from that species in that the leaves have an acumen (0.5 – )0.7 – 0.8(– 1.2) cm long vs 0.3 – 0.5 cm, secondary nerves are 7 – 10 (– 12) on each side of the midrib (vs 4 – 6 in *S. mpassa*), and that the stipules are 5 – 6.5( – 7) × 3 – 5(– 6) mm ( vs 2.5 mm long) and truncate (vs overtopped by an awn c. 1 mm long). See also Table 2.

**Table 2.** Characters separating *Sericanthe mpassa* from *Sericanthe onanae sp.nov.* Data for the first species taken from Sonké *et al*. (2012) and from the type herbarium specimen at K.

|  | *Sericanthe mpassa* | *Sericanthe onanae* sp. nov. |
| --- | --- | --- |
| Altitudinal range and habitat | c. 300 m alt. rainforest | 1200 – 1575 m alt. submontane forest |
| Habit | Small tree, height unknown. | Tree 8 – 25 m tall. |
| Leaf axillary buds | Conspicuously hairy, hairs c. 1 mm long, patent, dense, white | Hairs inconspicuous or glabrous |
| Abaxial leaf surface colour & surface | Green, matt | Golden brown, glossy |
| Indumentum of stems, stipules, abaxial leaf nerves when young | c. 0.5 mm long. caducous | c. 0.05 – 0.2 mm long, persistent |
| Stipular sheath length, overtopped by awn (length | Sheath 2.5 mm long, awn c. 1 mm long | Sheath $5 - 6.5(-7) \times 3 - 5(-6)$ mm long, awn absent. |
| Number of veins each side of the midrib | 4 – 6 | 7 – 10 (– 12) |
| Acumen length (cm) | 0.3 – 0.5 | (0.5 – )0.7 – 0.8(– 1.2) |
| Peduncle | Visible both above the proximal and below the distal calyculi | ±concealed by the calyculi |
| Corolla mouth hairs | Hairs absent or inconspicuous | Completely occluded by a shallow dome of bright yellow dense hairs, apart from a small central aperture |
| Style indumentum | Absent | Present |
| Geographic Range | Gabon | South West Region<br>Cameroon & Equatorial<br>Guinea, Bioko |

<u>Evergreen tree</u> 8 – 25 m tall. Leafy stems 20 – 30 cm long, unbranched, internodes (first 2 – 4) purple-black, terete, internodes (0.7 –)1.1 – 3.0(– 3.3) cm long, 2 – 4 mm diam., becoming roughly square in section, epidermis glossy gray, longitudinally furrowed, 4 – 6 mm diam. at flowering nodes; 2 – 4( – 5) pairs of leaves present per stem; glabrous (Fig. 2A) except the c.0.5 mm above the stipule insertion (completely concealed by the persistent stipules) where minutely white, simple pubescent, hairs 0.02 – 0.05 mm long. <u>Stipules</u> long-persistent, brown, leathery in proximal part, subcylindrical, 5 – 6.5( – 7) × 3 – 5(– 6) mm, completely sheathing, apex truncate, midrib ridged distally for c. 2 mm long (Fig. 2C&D), not overtopping the sheath as an awn; stipules of older internodes (4th – 5th or more from the apex) often splitting longitudinally into two halves; outer surface with sparse (c. 20% cover) simple white, appressed hairs c. 0.2 mm long, caducous in older stipules, inner surface glabrous apart from the basal c. 0.5 mm with dense silky hairs c. 1.8 – 2(– 4) mm long mixed with numerous dull red, winged colleters (Robbrecht 1987a), c. 2 x 0.05 mm secreting a translucent gum embedding the hairs and colleters together and adhering them in a layer to the inner surface of the stipule, completely covering it, and often exuding from the stipule onto the stem and expanding leaves giving them a glossy translucent coating; longest inner stipule hairs sometimes exserted from the sheath (Fig 2D). <u>Leaf-blades</u> papery, upper surface drying dark brown, lower surface drying glossy golden-brown, the abaxial midrib purple-brown; oblanceolate (elliptic in *Carvalho & Fernandez Casas* 4169-1), (6 – )11.2 – 13.8( – 15) × (3.8 – )5.3 – 6 (– 6.8) cm, apex rounded to truncate then abruptly triangular-acuminate, acumen (0.5 – )0.7 – 0.8(– 1.2) x (0.5 – )0.7 – 0.9(– 1.2) cm, base of blade acute, slightly asymmetric, shortly decurrent down the distal petiole; adaxial surface with midrib and secondary nerves sunken, glabrous; abaxial surface with midrib raised, ±brochidodromous (secondary nerves seemingly looping); domatia absent (Fig. 2B); bacterial galls/nodules probably absent (not detected) or at least very inconspicuous; secondary nerves orange or purple, 7 – 10(– 12) on each side, arising at 45 – 70 degrees from the midrib, curving upwards and gradually joining the nerve above through a branch, forming a weakly looping infra-marginal nerve; hairs minute, 0.08 – 0.2 mm long, white appressed, sparse (c. 5 – 10% cover Fig 2B), along the midrib, secondary nerves and margin abaxially, otherwise glabrous on both surfaces; tertiary nerves mostly highly inconspicuous (view with lense) barely raised, concolorous, scalariform, patent to midrib, 15 –20 between each pair of secondary nerves), sparsely branched, glabrous; quaternary nerves inconspicuous; margin slightly recurved and thickened. <u>Petiole</u> spreading, shallowly canaliculate (0.7 – )1.3 – 1.5( – 1.8) × 1.1 – 1.4 mm, inconspicuously hairy, hairs white 0.2 – 0.3 mm long, white appressed, about 20% cover along sides (Fig. 2C&D). <u>Inflorescences</u> on older stems several or many nodes below the leaves, supra-axillary or not, in both axils of a node (commonly a second or third superposed inflorescence per axil) (Fig. 2E), flowering/fruiting nodes per stem commonly (4 –) 10 –15 occurring on the previous seasons stems, from the (5 –) 9 – 24 th nodes from the apex, flowers 1 per inflorescence, inflorescence highly condensed; mature fruit and flower buds occurring at the same node. Calyculi 3 – 4 per flower, increasing in size distally, papery, ±cup-shaped, either truncate or with two foliar and two stipular awns attached below the margin, outer surfaces inconspicuously hairy, hairs c. 60 % cover, white, appressed, forward directed or randomly oriented, 0.05 – 0.1 mm long; inner surfaces glabrous but completely concealed by a layer of basally attached colleters and hairs, as the stipules (see above), longest hairs sometimes exserted (Fig 2F); axis completely concealed by calyculi in flower for all but 0.5 – 2 mm of its length (between the 1^st^ and 2^nd^ order calyculi, Fig. 1F), axis c. 0.9 mm diam., indumentum as outer calyculi surface; axis mostly visible in fruit (Fig. 1E), 1^st^ order (proximal) calyculus brown, papery, shortly cylindrical to cupular, 0.5 – 2 × 2 – 2.5 mm, awns if present 1.6 –2.2 mm long, robust, erect; 2^nd^ order calyculi (if present), 1.8 x 1.9 mm, margin usually truncate, or awns 0.2 – 1.2 mm; 3^rd^ order calyculi, 1.5 – 2.5 x 3 – 3.8 mm, usually with two long awns c. 2 mm long, and two short awns c. 0.2 mm long; 4^th^ order calyculus (2 – )2.5 – 3.8 x 3.8 – 4.2 mm, shortest and widest when fissured, long awns 1.4 – 1.9 mm long, short awns c. 0.5 mm long, concealing the ovary-hypanthium at anthesis. <u>Flowers</u> hermaphrodite, homostylous, 8-merous, 19 – 20 mm diam. c. 10 mm long (corolla and stamens). Ovary-hypanthium urceolate, c. 1.5 x 1.75 mm, concealed within distal calyculus, apex appressed hairy, otherwise glabrous. Calyx in bud ellipsoid, apex truncate - apiculate, not covering the corolla in bud, apiculae (visible only in bud) c. 0.5 mm long; calyx tube at anthesis subcylindrical, widest at apex, c. 2.8 x 2.5 mm; lacking longitudinal fissures after anthesis (Fig. 2F-H); calyx lobes pronounced post-anthesis, 8, shallowly rounded, lobes 0.6 – 2.1 mm, becoming inflexed as fruit develops (Fig. 2G); base (disc area) post-anthetically becoming longitudinally furrowed at apex; outer surface indumentum as calyculi; inner surface brown, glabrous apart from a few extremely sparse and inconspicuous, widely scattered slender white hairs 0.1 – 0.2 mm long; colleters not seen; raised longitudinal ribs 2 – 3 per lobe (Fig. 2H). Corolla apex exposed above calyx in bud, conical, apex mucronate; drying orange-brown, tube cylindrical, 3.75 – 4 × 2.25 – 3.5 mm, very slightly widest at mouth, outer surface glabrous, apart from 8 slender longitudinal lines alternating with the lobes; lines c. 0.15 mm wide, of bright white appressed hairs each 0.05 – 0.2 mm long, hair lines separated from each other by 0.4 mm; proximal 1.5 mm tune entirely glabrous; inner surface with proximal ¾ (3 – 3.5 mm) glabrous, hair band distal, c. 0.75 mm long, with extremely dense highly crinkled flat bright yellow hairs 1 (– 2) mm long, nearly occluding the mouth, and surrounding the bases of the staminal filaments. Corolla lobes 8, left-contorted in bud, spreading at anthesis and longer than tube, obovate, 9–10 × 5 – 7 mm, apex obtuse, adaxially glabrous, abaxially white puberulent with a slender glabrous midline, hairs c. 0.1 mm long, appressed. Stamens 8, anthers fully exserted, glabrous, anthers lanceolate-sagittate, 4 – 5 × 1.8 – 2.5 mm, connective conspicuous the entire length, drying dark brown, basifixed, apical connective appendage absent; filaments 1.5 – 2 x 0.6 mm, inserted at mouth of tube, slightly dorsiventrally flattened. Disc subcylindrical, 1 × 1.5 mm, apex shallowly domed, glabrous, accrescent in post-anthetic flowers and developing 3 steps (see Fig 2J). Style 9 – 10.5 × 0.4 – 0.6 mm (narrowest at base, widest at apex), exserted, glabrous at apex, otherwise with straggling white hairs c. 0.25 – 0.75 mm long; style arms 2, forming a V-shape, each ellipsoid, c.1.5 x 0.6 mm, stigmatic surfaces smooth (Fig. 3C). Ovary 2–locular, placenta attached towards apex, pendulous, elliptic, ovules 2 per locule, inserted laterally and partly immersed in the swollen placentae. Fruit ripening from green to yellow to orange when mature (*Ashworth* 337), subglobose or shortly ellipsoid, 10 – 15 x 10 – 12.5 mm, apex crowned by the accrescent cylindrical calyx c. 5 x 4 mm (Fig. 2E), calyx tube thickened, lobes papery, brown; disc slightly accrescent, c.5 mm wide (Fig 2J); base covered with the appressed, divided calyculus; peduncle slightly accrescent 8 x 1.5 mm, naked apart from persistent calyculi; fruit wall leathery-fleshy, outer surface rough, glabrous, about 1 mm thick, endocarp parchment-like, 1 – 4-seeded. Seeds ellipsoid (2-seeded fruit, slightly immature, *Onana* 1949) c. 10 –13 x 8 – 11 x 5 – 6 mm, testa glossy medium brown, hilum white, covering the top 25 – 30% of surface as a cap on the abaxial surface extending in a tapering line proximally, endosperm bright white, embryo erect (not transverse), directed distally, cylindric-conical c. 5 x 1.2 mm, cotyledons flat c. 2 x 1.2 mm.

**Fig. 2.**
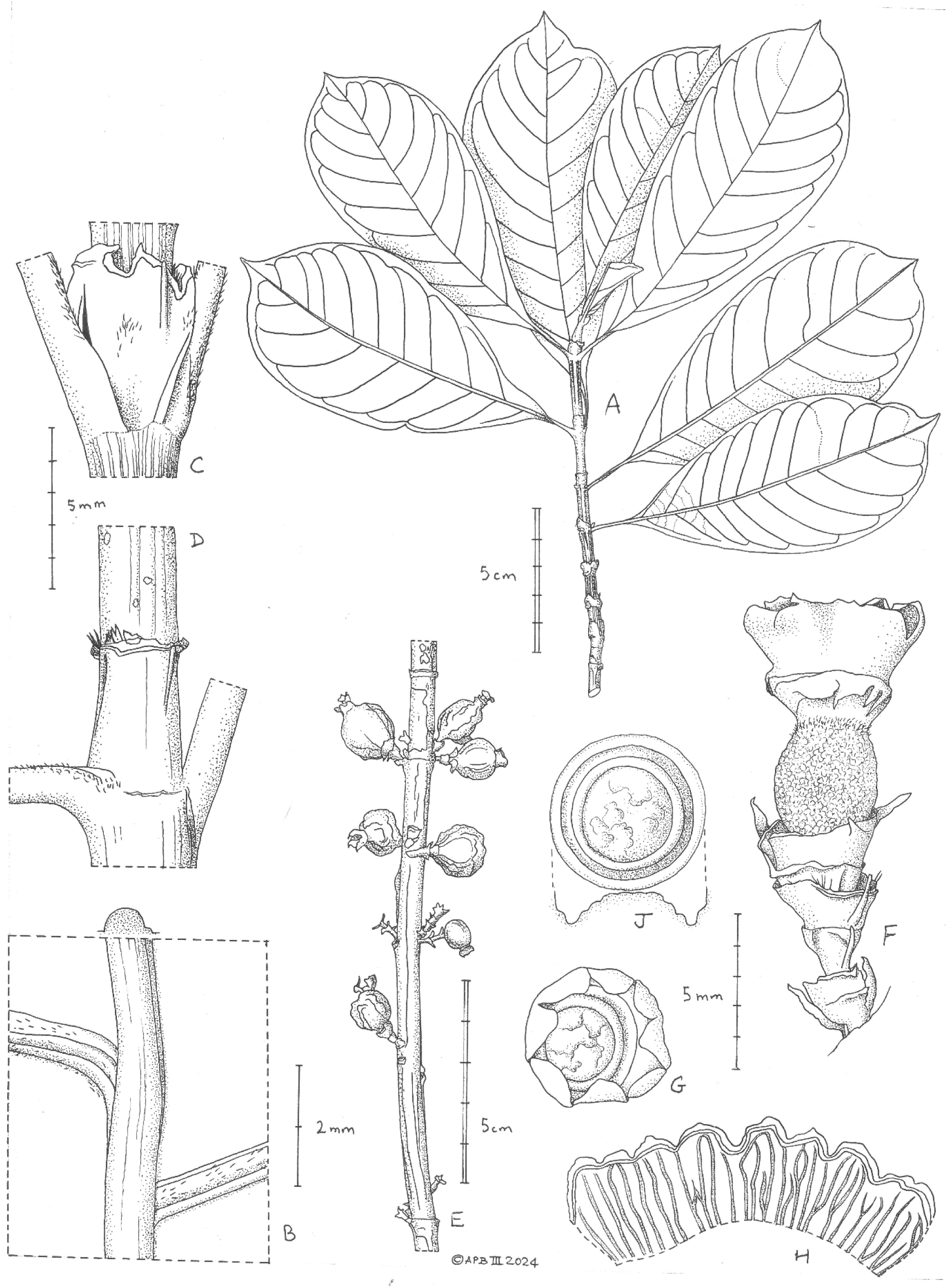
*Sericanthe onanae* **A** habit; **B** leaf abaxial surface showing midrib and two secondary nerve bases (domatia absent); **C & D** awnless sheath stipules (above damaged, from old internode); **E** infructescences on old leafless stem; **F** post anthetic flower/immature fruit, showing calyx-hypanthium and four calyculi, corolla, stamens and style, side view; **G** calyx from F view from above to show inflexed calyx lobes; **H** inner surface of calyx from G, opened and flat to show nervation; **J** view of disc from apex of mature fruit, with profile **A-C, E** from *Onana* 1949 (K); **D** from *Etuge* 3961 (K); **F-J** from *Ashworth* 337 (K). Drawn by ANDREW BROWN.

**CAMEROON. SOUTH WEST REGION. Banyang Mbo Wildlife Reserve**. Ebamut, 1200 m alt., fr. 8 Nov. 2001, *Onana* 1949 with Nana, Ghogue, Etuge, Ambrose, Okong, Kongor, Fobia (K holo.; iso. BR, YA); **Bakossi Mts**. Kodmin, Muawhonzum Trail, 1500 m. fr. 17 Jan. 1998, *Etuge* 3961 with Clement, Innocent, Victor, Duncan, Lynne (K, MO, YA); **Kagwene.** Cocoyam farm at bottom of Alumfa path, 1575 m, old fl., fr. 17 June 2009, *Ashworth* 337 with Ndo, John; Okon, Felix; Anya, Sylvester (BR, K, MO, YA); **EQUATORIAL GUINEA. BIOKO.** Metadata mislaid, possibly collected (data from Carvalho 4188): Moka to Ureca 1146 m alt., fl. 30 Nov. 1999, *Carvalho & Fernandez Casas* 4169-1 (K 2 sheets).

**HABITAT & ECOLOGY**. Submontane forest; 1200 – 1575 m alt.

**DISTRIBUTION**. Cameroon, Equatorial Guinea (Bioko).

**ETYMOLOGY.** Named (meaning Onana’s silky flower) for Jean Michel Onana, former Director of the Cameroon National Herbarium, author of the Cameroon checklist of Vascular Plants (Onana 2011), co-author of the Cameroon Red Data book of Plants (Onana & Cheek 2011) and champion of the conservation of threatened plant species in Cameroon. He is commemorated by four other species: *Ledermanniella onanae* Cheek (Podostemaceae), *Diospyros onanae* Gosline (Ebenaceae), *Psychotria onanae* O.Lachenaud (Rubiaceae) and *Deinbollia onanae* Cheek (Sapindaceae). For further information see Cheek *et al*. (2022a).

**CONSERVATION STATUS.** At each of the four sites from which this species has been recorded by a herbarium specimen there are threats of habitat clearance from small holder agriculture. In fact, on one specimen the location is stated as “cocoyam farm under canopy” (*Ashworth* 337, K from Kagwene). This site is apparently outside a protected area and the note indicates that the understorey of the forest beneath the tree had been cleared for this crop (*Colocasia esculenta* (L.) Schott) which would limit or halt the possibility of regeneration from seedlings of the species, even if the forest canopy remains mainly intact. This same threat of small holder agricultural clearance occurs at the site of *Etuge* 3961 which is <700 m distant from the village of Kodmin, the main settlement at altitude in the Bakossi Mts, and outside the National Park of that name. The site of the type specimen is near the village of Ebamut which is inside the Banyang Mbo Wildlife Sanctuary, but nonetheless agriculture from the village is currently a source of threat. Ebamut village grew slightly between 2010 and 2021 (images on Google Earth Pro, time slider function). The Bioko site is estimated to be likely within a kilometre or two of the town of Moka so again, small holder agriculture is a threat. All three sites in Cameroon are currently inaccessible due to the ongoing armed struggle for independence by the anglophone inhabitants of South West and North West Regions, Cameroon that began in December 2016. Using the IUCN required 2 km x 2 km grid cells we calculate the area of occupancy as 16 km^2^. The extent of occurrence was calculated as 6307 km^2^ which qualifies for the Vulnerable level of threat. However, the AOO, together with the threats stated, allows us to make here a provisional assessment of Endangered EN B2ab(iii) for *Sericanthe onanae*.

It is possible that the species may be found at other sites, but numerous surveys have already been conducted in the range and surrounding area of this species (detailed under the previous species, *Sericanthe etugei*).

The first step to help protect this species should be a sensitisation programme with the local communities concerned to enable understanding of the importance of conservation of this species and to enable species identification and protection. Development of an action plan to protect the species is recommended. Since seed seems to be abundantly produced (three of the four specimens have numerous fruits) propagation from seed in local nurseries is a possibility.

**NOTES.** The stipules of *Sericanthe onanae* are remarkable for being completely sheathing and lacking both a distal limb and an awn that overtops the sheath, because the thickened ridge-like midrib hardly develops a distal extension. Several other species of the genus also have such a sheathing stipule but these have a 1 – 2 mm long awn (*S. jacfelicis*, *S. gabunensis*), or in the case of *S. mpassa* c. 1 mm long (Robbrecht 1978; Sonké et al. 2012). However, *S. etugei* approaches the state of *S. onanae* since the awn is extremely reduced, only c. 0.25 mm long (this paper, above). In some species, such as both *S. onanae* and *S. etugei,* the sheath usually splits into two halves after about the 4^th^ node from the apex (so the stipule appears to have two limbs and a very short or no sheath) as the stem diam. increases in diameter. In *S. etuge* the awn can appear to elongate as the stipule ages, as the membranous sheath fragments and falls from the midrib.

The quantity of exudate from the colleters of the distal stipules seems similar to that in *S. mpassa* and more copious than in other species, although no detailed study has been made. Sometimes beads of exudate are seen at the edge of the stipule.

The urceolate ovary-hypanthium-calyx base, seen as a broad, longitudinally furrowed band of wider diameter than the top of the ovary and base of the calyx in the post-anthetic flower (Fig. 2 F), appears similar to that in *S. mpassa* but is otherwise unusual in the genus and appears correlated with the unusually large (long and broad) development of the floral disc in this species. However, it may be that this structure occurs in further species for which suitable material for viewing is unavailable.

The alternating longitudinal glabrous and hairy bands seen on the external corolla tube are also remarkable in being unreported in other species of the genus. The bands are variable in width, and were present in all but one of c. 30 well preserved flowers examined. However, they may be a monstrosity.

The bright yellow (dried state) and very long, crinkled and dense hairs of the corolla throat form a shallow dome c. 5 mm wide and c.0.5 mm high at the top of the corolla, from inside the margins of which the staminal filaments emerge, and in the middle of which is a small aperture, c. 0.3 mm wide, through which the style is placed.

*Sericanthe onanae* is unusual in the genus being recorded as a tree 25 m tall (*Ashworth* 337, K). It exceeds maximal heights given for other species in the genus, e.g. *S. toupetou* (Aubrév. & Pellegr.) Robbr. at 20 m, and *S. roseoides* (De Wilde & Th. Dur.) Robbr. at 15 m (Robbrecht 1978).

This taxon was first named as *Sericanthe* sp. B (Cheek et al. 2004) based on *Etuge* 3961 from the Bakossi Mts near Kodmin, which is in fruit. *Ashworth* 337 (fruit and post-anthetic flowers) specimen and that of *Onana* 1949 (flower buds and fruit) were collected subsequently.

The specimen from Bioko (*Carvalho & Fernandez Casas* 4169-1, K) was only found in the closing stages of this paper. The metadata for this specimen was mislaid and so we have extrapolated it from *Carvalho* 4188, about 20 numbers distant, in order to have an approximation of the locality and date of collection. However this is uncertain. For this reason, and because no duplicates have been found, this specimen has not been selected as type even though it is otherwise of excellent quality and is the only specimen with open flowers. It had been determined as *Sericanthe jacfelicis vel sp. aff.* by Davis in 2003. It is not referred to in Sonké et al. (2012), nor indeed are any specimens of *Sericanthe* from Bioko noted in that or any other publication that we have found. Searching gbif.org of the genus from Bioko shows four records, regarding 2 unique collection events, each a single species, identified as *S. auriculata* (*Luke* 13258, K) and *S. jacfelicis* (*Luke and Fermin* 12105 EA, MA, MO all n.v.). The last species was collected in lowland forest and therefore is unlikely to be additional material of the new species. Therefore Bioko has three species of *Sericanthe.* Enquiries at MA for *Carvalho & Fernandez Casas* 4169-1 failed to find duplicates there.

Morphologically the species seems vegetatively uniform. *Carvalho & Fernandez Casas* 4169-1 is slightly discordant from the other specimens in that the leaf blades are elliptic and not obovate, but otherwise it appears concordant.

We speculate that seeds of this species might be dispersed by primates such as gorilla and chimpanzees, since the orange leathery-coated fruits common in the genus contain large seeds each partly or mostly enveloped in a white fleshy placental-arilloidal layer that is likely edible.

### Submontane species discovery in the Cross Sanaga Interval

*Sericanthe etugei* and *S. onanae* are further additions to the very large number of submontane species (altitudinal range approximately 800 to 2000 m in Cameroon, Cheek *et al*. 2004) that occur in the Cameroon Highlands of the Cross Sanaga Interval. In recent decades these numbers have been greatly amplified. The species vary from terrestrial herbs: Champluvier & Darbyshire (2009); Cheek & Xanthos (2012); Cheek *et al*. (2023a); Darbyshire *et al*. (2011); Maas-van de Kamer *et al*. (2016); Pollard (2024), to rheophytes: Cheek (2003); Cheek & Ameka (2008); Cheek *et al*. (2017; 2022b); epiphytes: Cheek & Csiba (2002a); Cheek & Fischer (1999); Muasya *et al*. (2010), shrubs: Cheek (2002); Cheek & Csiba (2000; 2002b); Cheek & Etuge (2009a); Cheek & Sonké (2005); Cheek *et al*. (2008; 2009; 2020a); Gosline & Cheek (1998); Hoffmann & Cheek (2003); Sonké & Bridson (2001); Stoffelen *et al*. (1997); Stone *et al*. (2024); to trees Cheek (2017); Cheek & Etuge (2009b); Cheek & Ngolan (2007); Cheek & Onana (2021); Cheek *et al*. (2002a; 2002b; 2017; 2018c; 2020b; 2023; 2024); Kenfack *et al*. (2003); Lachenaud *et al*. (2024); Onana & Chevillotte (2015); Sonké (2000); lianas e.g. Hoekstra *et al*. (2016; 2021); Cheek & Onana (2024), and mycoheterotrophs: Cheek *et al*. (2003; 2018d; 2019; 2023c).

## Conclusion

It is critical to detect, delimit and formally name species as new to science as soon as possible now, since until they are scientifically published they are effectively invisible to science. Only when species have a scientific name can their inclusion on the IUCN Red List be facilitated (Cheek *et al*. 2020c). Most (77%) species named as new to science in 2020 were already threatened with extinction (Brown *et al*. 2023). Many new species to science have evaded detection until today because they have small ranges which have remained unsurveyed for plants until recently, as was the case with the two new species of *Sericanthe* described in this paper.

Cameroon has the highest number of globally extinct plant species of all countries in continental tropical Africa (Humphreys *et al*. 2019). The extinction of species such as *Oxygyne triandra* Schltr. (Thismiaceae, Cheek *et al*. 2018d) and *Afrothismia pachyantha* Schltr. (Afrothismiaceae, Cheek & Williams 1999; Cheek *et al*. 2019; Cheek *et al*. 2023c) are well known examples. Examples of apparently extinct tree species include *Vepris bali* Cheek (Rutaceae, Cheek *et al*. 2018c) and *Vepris montisbambutensis* Onana (Onana & Chevillotte 2015). However, another 127 potentially globally extinct Cameroon species are documented (Onana in Murphy *et al*. 2023: 18 – 22). Cameroon also has a vast number of highly threatened species (Onana & Cheek 2011), including the highest number (414) of threatened tree species of all tropical African countries (BGCI 2021). These will be further increased by the addition of the two new species of tree published in this paper.

If further global extinction of plant species is to be avoided, effective conservation prioritisation, backed up by investment in protection of habitat, ideally through reinforcement and support for local communities who often effectively own and manage the areas concerned, is crucial. Important Plant Areas (IPAs) programmes, often known in the tropics as TIPAs (Darbyshire *et al*. 2017; Murphy *et al*. 2023) offer the means to prioritise areas for conservation based on the inclusion of highly threatened plant species, among other criteria. Such measures are vital if further species extinctions are to be avoided of narrowly endemic, highly localised species such as *Sericanthe etugei* and *S. onanae*.

## Acknowledgements

We thank Diane Bridson and Elmar Robbrecht for reviewing an earlier version of this manuscript. The botanical surveys in Cameroon which resulted in this paper were mainly supported by Earthwatch Europe (1993 – 2005) and by the Darwin Initiative of the UK Government through the Plant Conservation of Western Cameroon, and the Red Data Book of Cameroon projects, both led by RBG, Kew, working with the IRAD-National Herbarium of Cameroon. The visit to Kagwene Gorilla Sanctuary was supported by Cameroon Wildlife Conservation Society (CWCS), and that to Ebo by the Ebo Forest Research Project (with Zoological Society of San Diego’s Institute for Conservation Research). Gaston Achoundong, former head of the National Herbarium of Cameroon (YA) and his successors including Barthelemy Tchiengué are thanked for their collaboration and support over the years. Our thanks also go to Jeannette Mapi-Sonké and the authorities of the University of Yaoundé I for their support to the last author’s (BS) research activities.

